# Basic-peptide depletion and precursor masking in bottom-up proteomics

**DOI:** 10.64898/2026.11.10.765051

**Authors:** Shoab Saadat, Michael R. Barnes, Goran Mohammad, Syeda Naveen Batool Naqvi, Christoph Thiemermann, Magdi Yaqoob, Vahitha Abdul-Salam

## Abstract

Biomarker proteomics studies report protein identities at the canonical precursor level, yet the underlying mass-spectrometry evidence consists of peptides whose physicochemical properties may diverge from the annotated protein. We reanalyzed peptide-level evidence from public datasets with a proteoform-aware pipeline that maps detected peptides to canonical sequences under a five-level confidence taxonomy, computes sequence-derived physicochemical features for precursors, peptides, and detected regions, and tests class-composition bias against digestion-matched backgrounds. Across ten public studies spanning five result-file formats and 70,020 peptide-evidence rows, detected peptides are depleted in the basic isoelectric-point class relative to their own proteins’ theoretical tryptic peptides (fixed-effect odds ratio 0.45, 95% CI 0.44–0.46; leave-one-study-out envelope 0.44–0.49). The depletion is absent among fully tryptic peptides (OR 1.06) and concentrates in missed-cleavage peptides (OR 0.62–0.57), is invariant to detected-abundance stratification, and retains its direction under three independent pI-calculator models. Case studies reveal both directions of precursor-centric masking. Two orthogonal validations ground the framework: a digestion-free synthetic-peptide analysis (basic-class odds ratio 0.73, 95% CI 0.71–0.76 under standard DDA) and a paired top-down benchmark, in which detected regions recover top-down sequence space (90% of spans overlapped) but do not delimit proteoform boundaries (8% containment). Pipeline, data, and code are public.

## Introduction

Bottom-up proteomics is the dominant workflow for plasma and serum biomarker discovery: proteins are digested into peptides, tandem spectra identify the peptides, and protein identities are inferred from them^1,2^. The physicochemical properties of the detected material are therefore peptide-level, yet biomarker candidates are reported as canonical precursor proteins with precursor-level annotations^3^.

When a protein is identified by peptides that cluster in one region of its sequence, the detected evidence represents that region—not the full precursor^4^—and precursor-level reporting inherits the canonical annotation’s isoelectric point and molecular weight even when the region’s own differ. Peptide-centric approaches have recognised this gap for individual datasets^5,6^ and correlation-based methods can recover proteoform-group structure^7^, but no cross-study framework has quantified precursor-versus-detected divergence at repository scale.

The proteoform concept^8,9^ provides the vocabulary (Fig. 1a). A reported biomarker “protein” may represent a canonical precursor, a protein group, a single peptide, a detected region, or a candidate proteoform; each level carries different inference confidence. If precursor-centric annotations systematically misrepresent the physicochemical space of detected evidence, the bias should be measurable against a digestion-matched background—the theoretical tryptic peptides of the same parent proteins—so that shared precursors isolate detection bias from protein-level composition.

**Figure 1.**
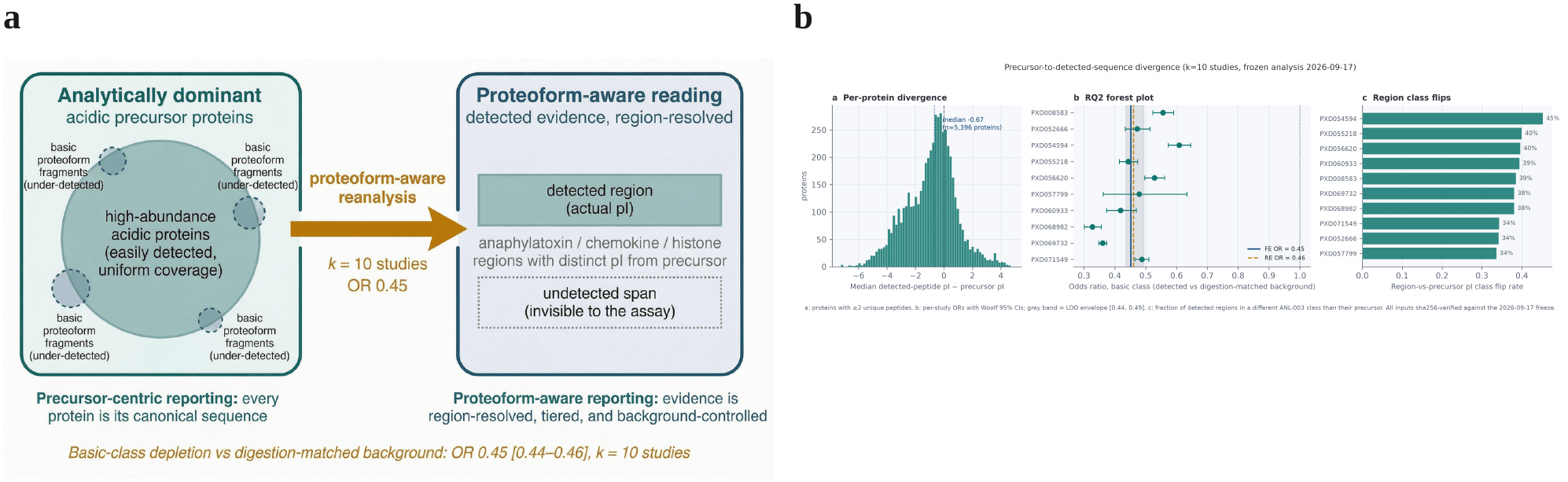
| Conceptual basis and the headline divergence and depletion result. **a**, The Yin and Yang of biomarker proteomics: under precursor-centric reporting (left), high-abundance acidic proteins dominate detection with uniform coverage while basic proteoform fragments are under-detected; the proteoform-aware reading (right) resolves detected evidence to its actual regions with their own physicochemistry. **b**, Precursor-to-detected-sequence divergence and basic-class depletion across the frozen ten-study cohort. Left: per-protein divergence (median detected-peptide pI minus precursor pI) for 5,396 proteins with ≥2 unique peptides; median −0.671 pI units (blue dotted line) vs digestion-matched permutation-null centre +0.116 (p = 0.0005). Middle: forest plot of basic-class depletion (RQ2) with per-study odds ratios and Woolf 95% CIs; fixed-effect 0.452; random-effects 0.461; leave-one-study-out envelope. Right: fraction of detected regions in a different pI class than their precursor, per study. All inputs sha256-verified against the 2026-09-17 freeze.

Sample-preparation workflows can under-detect small and basic proteins^10^; peptide detectability depends on sequence, instrument, and acquisition mode^11^; and plasma and serum span a wide dynamic range^12,13^. Complement anaphylatoxins, chemokines, and histone-derived fragments anchor the biological case studies^14,15,16^.

Our framework addresses three questions: whether detected peptides diverge in physicochemical space from their canonical precursors beyond what random sampling of the same proteins’ theoretical tryptic peptides would predict (RQ1); whether detected peptides exhibit a systematic class-composition bias—depletion of basic-class peptides—relative to those digestion-matched backgrounds (RQ2); and, in an exploratory analysis, whether disease-annotated functional regions show differential basic-class enrichment (RQ3).

This study addresses two gaps: no prior work has quantified precursor-versus-detected physicochemical divergence against digestion-matched backgrounds across studies, file formats, and disease contexts; and no reporting taxonomy distinguishes peptide-level from region-level from proteoform-level evidence in biomarker databases. We contribute:

- A proteoform-aware reanalysis pipeline (five parser families, join-quality gate, benchmarked physicochemical engine, five-level confidence taxonomy) that resolves bottom-up evidence to the sequence regions actually observed, quantifies their divergence from canonical annotations, and exposes acquisition-dependent detection bias.
- A cross-study quantification of basic-class depletion relative to digestion-matched backgrounds, with calculator-sensitivity validation, covariate stratification, and leave-one-study-out stability.
- A mechanistic localization of the depletion to missed-cleavage peptides, with charge-state evidence linking uncut basic residues to elevated precursor charge.
- Six case studies demonstrating both directions of precursor-centric masking, anchored by a methodological null control.
- Orthogonal validation on two independent resources under metric contracts frozen before computation: digestion-free detectability across 214,336 synthetic peptides, and benchmarking against paired top-down proteomics, which supports sequence-space recovery (90% overlap) but not boundary delimitation (8% containment).

## Results

### Cohort and pipeline overview

We screened 135 public biomarker proteomics studies from the PRIDE Archive^26^ and selected ten for full extraction under a written rubric scoring parser-family balance, file accessibility, sample-type fit, metadata quality, and fractionation-evidence availability (Methods; Fig. 2). The selected cohort spans five result-file formats: MaxQuant peptides and PSM tables (three studies), DIA-NN precursor reports (three), Spectronaut facility peptide quantification (one), Progenesis CSV (one), and mzIdentML (one), plus a site-specific glycopeptide workflow (one). Seven datasets derive exclusively from plasma or serum; three include non-plasma material (FFPE intervertebral-disc tissue, FFPE colorectal-tumour tissue paired with plasma-derived extracellular vesicles, and cerebrospinal-fluid extracellular vesicles). Full extraction yielded 70,020 peptide-evidence rows covering 34,784 unique peptide sequences and 5,777 proteins resolved against UniProt^27^. The median per-parser-family pI ranged from 5.28 to 5.97, confirming cross-format convergence of the physicochemical engine (Supplementary Fig. S1).

**Figure 2.**
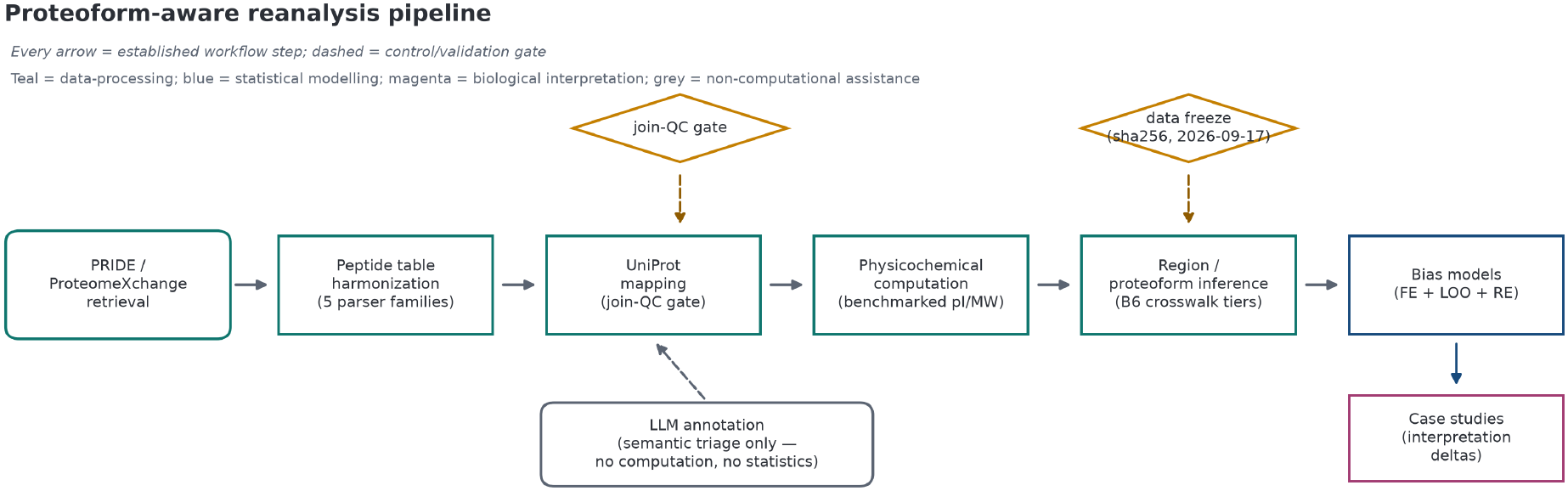
| Proteoform-aware reanalysis pipeline. Teal rectangles = data-processing steps from repository retrieval through region inference; blue = statistical modelling; magenta = case studies; diamonds (amber, dashed) = control and validation gates (join-quality gate and data freeze); LLM annotation (grey, dashed) = semantic triage only. The pipeline operates on ten public studies and quantifies basic-class under-representation at OR 0.45 [0.44, 0.46] vs digestion-matched backgrounds.

### Detected peptide evidence diverges from precursor annotations (RQ1)

For each protein with at least two unique detected peptides (n = 5,396), we computed the median difference between detected-peptide pI and the canonical precursor pI. The median per-protein divergence is −0.671 pI units—more acidic than a digestion-matched permutation null of +0.116 (2,000 resamples per protein at matched peptide counts; two-sided p = 0.0005; Fig. 1b). This confirms that the detected evidence is systematically more acidic than random sampling of the same proteins’ theoretical tryptic peptides would predict. Overall, 35.1% of detected peptides fall in a different pI class than their precursors under pre-specified thresholds (acidic < 7.0, near-neutral 7.0–7.5, basic ≥ 7.5).

### Systematic depletion of basic-class peptides (RQ2)

Compared with digestion-matched theoretical tryptic peptides from the same parent proteins, detected peptides are depleted in the basic pI class in every study individually (per-study odds ratios 0.33–0.61; Fig. 1b). The inverse-variance fixed-effect summary odds ratio is 0.452 (95% CI 0.443–0.461); the leave-one-study-out envelope is 0.436–0.494, confirming that no single study drives the estimate. A DerSimonian–Laird random-effects summary, reported because the cohort reached the pre-specified threshold of ten studies, is 0.461 (p = 7 × 10^−30^; I² = 97.6%, reflecting variation in magnitude, not direction). Correspondingly, detected peptides are over-represented in the acidic class (Fig. 3a–b).

**Figure 3.**
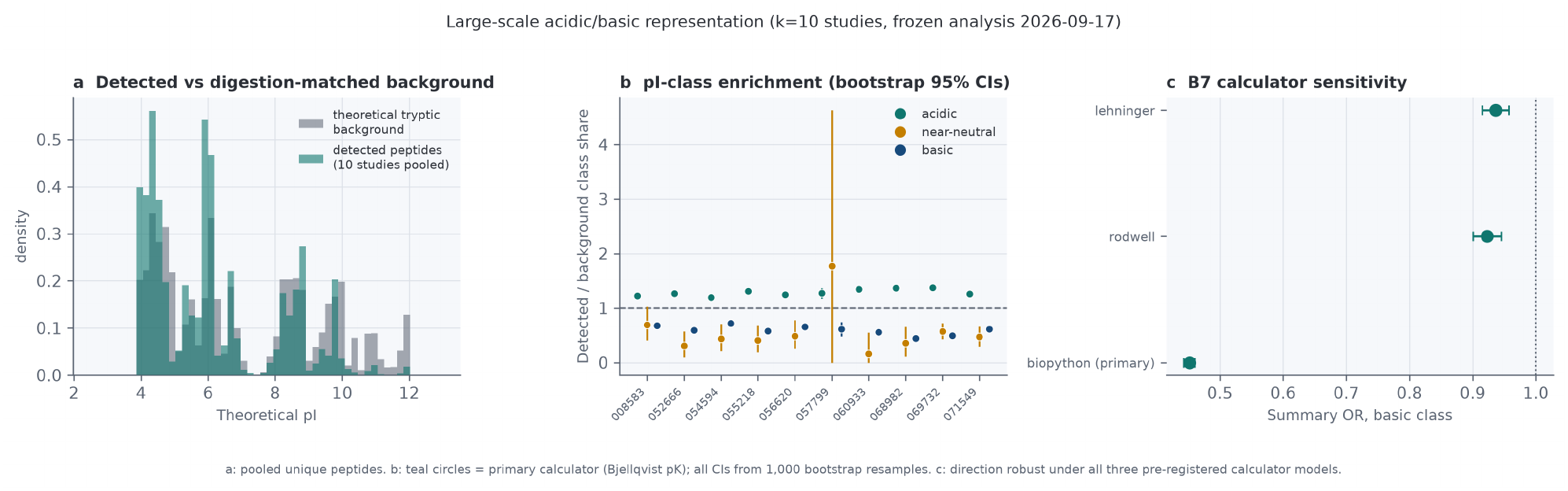
| Large-scale acidic/basic representation. **a**, Pooled unique detected peptides (teal) vs digestion-matched theoretical tryptic background (grey). **b**, pI-class enrichment ratios per study with bootstrap 95% CIs. **c**, B7 calculator sensitivity: summary basic-class ORs under three pre-specified pK models (Bjellqvist, Rodwell, Lehninger)—direction robust, magnitude calculator-dependent.

The conclusion’s direction is robust to calculator choice. Under the primary Bjellqvist pK model, the basic-class summary OR is 0.45; under two alternative pre-specified pK models (Rodwell/EMBOSS lineage and Lehninger), it attenuates to 0.92 and 0.94 respectively, with all confidence intervals below 1 (Fig. 3c). The attenuation arises from the known instability of the near-neutral transition zone (pI 7.0–7.5), where the three pK sets disagree most on the protonation state of histidine and terminal residues. The per-protein divergence histogram (Fig. 1b) provides calculator-independent evidence: the median shift of −0.67 pI units is a continuous-distribution property, not a threshold-classification artifact.

### The missed-cleavage locus and covariate stratification

Stratification by missed-cleavage count reveals that the basic-class depletion is absent among fully tryptic peptides (mc0 OR 1.06, 95% CI 1.03–1.09) and concentrates in peptides with one (mc1 OR 0.62) or two or more (mc2+ OR 0.57) missed cleavages (Extended Data Fig. 1). Charge-state analysis of the reportable subset (33,055 peptides from three studies; Supplementary Information) confirms that missed-cleavage peptides are observed at charge states ≥3 in 80.0% of cases versus 25.9% for fully tryptic peptides (OR 11.46), providing the analytical link between uncut basic residues and elevated precursor charge. The depletion persists across all peptide-length quartiles (weakest in the longest) and is invariant to detected-abundance stratification (below-median OR 0.407 vs above-median 0.405; Pearson r of log-intensity with pI = −0.02, n = 52,848), confirming that dynamic range does not explain the effect.

### Disease-relevant region classes show differential basic-class enrichment (RQ3, exploratory)

Among detected regions annotated with disease-relevant class flags (complement anaphylatoxin, chemokine, histone), Fisher exact tests with Benjamini–Hochberg correction show strong basic-class enrichment for histone-region (OR 7.7, q = 1.3 × 10^−9^), histone-tail-candidate (OR 44.5, q = 3.3 × 10^−9^), and chemokine-region classes (OR 4.9, q = 6 × 10^−6^), while complement-anaphylatoxin regions do not reach significance at this cohort size (OR 1.7, q = 0.112). These findings are exploratory and reported with their correction-adjusted q-values.

### Case studies reveal both directions of precursor-centric masking

Six case studies, selected by a pre-stated rubric from the frozen outputs, demonstrate the two components of the thesis (Fig. 4).

**Figure 4.**
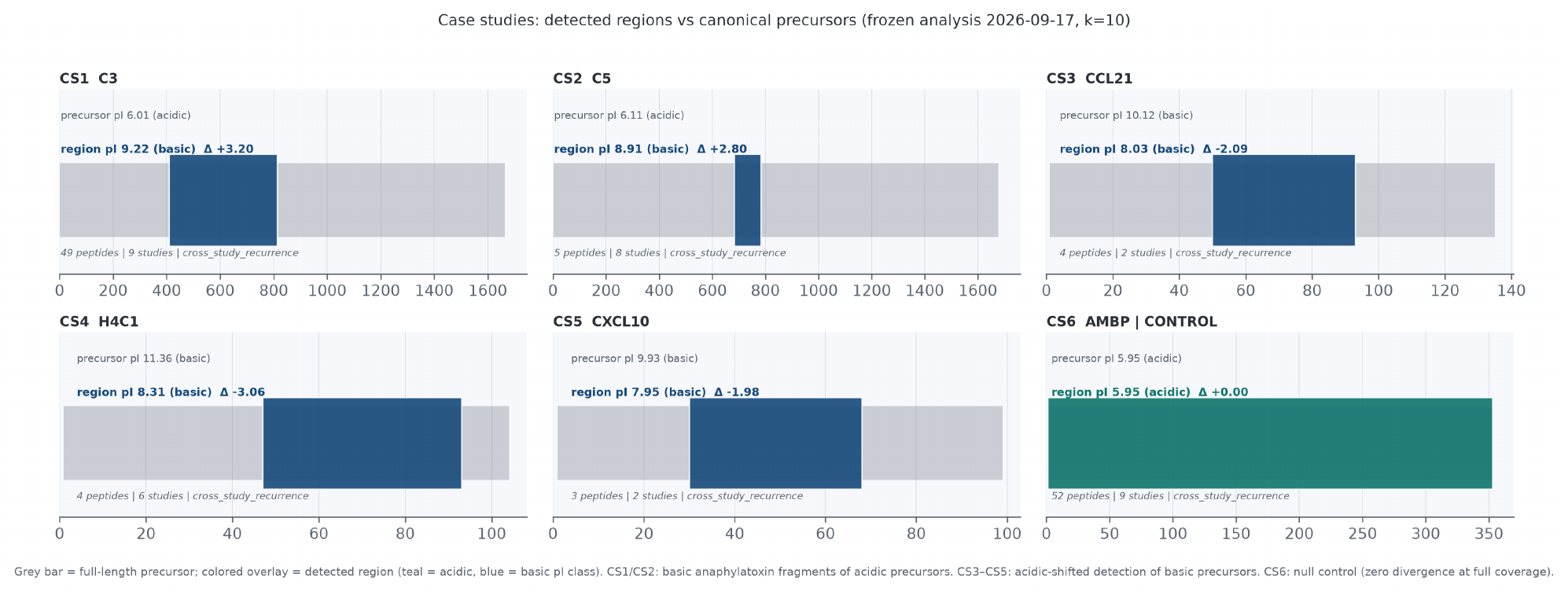
| Case studies: detected regions vs canonical precursors. Grey bar = full-length precursor; colored overlay = detected region (teal = acidic, blue = basic pI class). **CS1/CS2**: complement C3 and C5 anaphylatoxin regions—basic fragments (+3.2 and +2.8 pI units) of acidic precursors. **CS3–CS5**: chemokines and histone H4 tail—acidic-shifted detection of basic precursors. **CS6**: AMBP null control (zero divergence at full coverage). Peptide counts, study recurrence, and confidence tiers annotated per panel.

#### Counter-direction: reporting bias

Complement C3 (canonical precursor pI 6.02) was detected via 49 peptides clustering in the C3a anaphylatoxin domain (residues 672–748), whose minimal sequence footprint region has pI 9.22—a divergence of +3.20 units (nine-study recurrence). Complement C5 shows the same pattern in two independent studies (precursor pI 6.11; anaphylatoxin region pI 8.91; divergence +2.80). In both cases, the mass spectrometer successfully detected a basic inflammatory fragment; the standard reporting pipeline collapsed those peptides into the parent accession, labelling the finding as an acidic protein. Downstream biomarker registries indexed these as acidic species, rendering the basic fragment invisible to literature searches for basic biomarkers.

#### Forward direction: detection bias

Histone H4 (precursor pI 11.36) was detected via four peptides in the N-terminal tail region (residues 47–92; region pI 8.31; divergence −3.06; six-study recurrence). Chemokines CCL21 (10.13 → 8.03) and CXCL10 (9.93 → 7.95) show the same acidic shift. In these cases, the detected material is basic but measurably less so than the precursor annotation implies—evidence that the detection pipeline samples the less-basic portions of basic mediators.

#### Methodological null control

AMBP (alpha-1-microglobulin/bikunin precursor) was detected by 52 peptides across nine studies with uniform sequence coverage. Its detected-region pI equals the precursor pI to three decimal places (divergence 0.000). This demonstrates that the pipeline outputs zero divergence when coverage is uniform, confirming that the contrast cases reflect non-uniform detection, not computational artifacts.

A post-freeze, exploratory probe of the harmonized evidence found one clear splice-isoform case. In MASP1 (PXD055218, DIA), four Tier A peptides map only to the annotated MASP-3 isoform, within a 294-residue isoform-specific segment, alongside ten canonical peptides (Extended Data Fig. 8); a canonical-only label would conceal the species detected. Single study; no replication.

### Orthogonal validation: observed regions recover sequence space, not proteoform boundaries

Two pre-specified tests (metrics frozen before computation) benchmark the pipeline outside its discovery cohort. First, the paired top-down/bottom-up plasma-corona dataset of Sadeghi et al.^21^: harmonizing the published bottom-up PSM table (433,453 PSMs, 12 samples) through the standard evidence schema (accession resolution 99.8%, exact mapping 99.4%, join-quality gate passed) and running region inference unchanged, then benchmarking against 3,671 top-down proteoform spans, yields an honest split. Coverage is strong: 90.4% of top-down spans on the 237 evaluable proteins are overlapped by at least one detected region (protein-level bootstrap 95% CI 0.78–0.97), and the median top-down span is fully covered by the union of overlapping regions. Proteoform-boundary specificity, however, fails the pre-specified criterion: at the 0.80 containment rule, 41 of 510 evaluable regions (8.0%; 95% CI 5.7–11.0%) qualify, well below the 70% full-validation threshold, and the frozen adequacy rule therefore flags the published 25-residue stitching gap for re-examination (the precision-maximizing grid point, g = 5, reaches only 11.0%, so no gap parameter satisfies the criterion — the ceiling reflects bottom-up granularity, not the choice of 25; Extended Data Fig. 5). Where regions do qualify, their boundaries snap to top-down termini (median start offset 0 residues, 95% CI 0–2; median end offset −2, 95% CI −5 to 0; Extended Data Fig. 6), and the nested-span nuance of the as-published stitching rule alters only 18 of 15,640 regions (0.12%). The 41 concordant regions (39 proteins) populate the confidence taxonomy’s top tier as *partially validated*—plausible molecular boundaries, not complete proteoform inventories.

Second, a digestion-free test on 214,336 synthetic peptides from ProteomeTools^23,24^, where every peptide is present in its pool by construction: under standard DDA (222 runs, 158 pools), the basic isoelectric-point class remains under-detected relative to the acidic class (odds ratio 0.734, 95% CI 0.713–0.755; one-sided Fisher *p* = 9 × 10^−9^^9^). The deficit is specific to standard DDA—ETD shows none (OR 1.01) and multi-fragmentation acquisitions over-detect the basic class (1.16–1.19) —and is robust across the three pK models (0.727–0.734) and all length quartiles (0.64–0.75; Extended Data Fig. 7). Because digestion and matrix are absent by design, this confirms the basic-class depletion as a measurement phenomenon; the fully tryptic panel cannot test the missed-cleavage arm (stated as a limitation).

## Discussion

This work quantifies two connected biases in public biomarker proteomics. First, detected peptides diverge from canonical precursor annotations by a median of −0.67 pI units—more acidic than digestion-matched random sampling predicts—establishing that precursor-level reporting systematically misrepresents the physicochemical space of the actual evidence. Second, detected peptides are depleted in the basic pI class relative to their own proteins’ theoretical tryptic peptides in every study individually, a direction robust to calculator choice and to leave-one-study-out sensitivity. Together, these findings show that canonical protein-level labels conceal which sequence regions were observed, and that a region-resolved reading—validated for sequence-space recovery but not proteoform-boundary delimitation—exposes acquisition-dependent detection bias.

The missed-cleavage locus provides the mechanistic anchor. Fully tryptic peptides show no basic-class depletion (OR 1.06); the effect lives entirely in peptides with missed cleavages (OR 0.62– 0.57). Charge-state analysis confirms the physical link: missed-cleavage peptides carry z ≥ 3 in 80% of observations (vs 26% for fully tryptic), consistent with uncut lysine and arginine residues elevating precursor charge. Highly charged missed-cleavage peptides experience charge competition in electrospray ionization and generate complex higher-charge tandem spectra that database search engines historically score with lower confidence^11,17^. The depletion is therefore a digestion-and-detection phenomenon, not an intrinsic property of the proteome.

Several considerations bound the interpretation. The isoelectric point alone does not reliably predict electrospray response^17^; hydrophobicity and gas-phase proton affinity co-detect. A five-workflow cerebrospinal-fluid benchmark found no systematic pI bias for any method^18^; our plasma-and-serum-focused cohort may differ in matrix composition, or the digestion-matched background may be more sensitive to class-level effects than platform-level comparisons. Platform dependence is real^19^. We address these constraints by reporting three-calculator sensitivity (direction robust; magnitude attenuates from 0.45 to 0.92–0.94 outside the Bjellqvist pK family—a consequence of near-neutral transition-zone instability, not a failure of the approach), by providing the continuous divergence histogram (Fig. 1b) as calculator-independent evidence, and by stratifying by acquisition mode (DIA-only OR 0.438, k = 4; DDA-only OR 0.479, k = 6; per-study estimates in Fig. 1b) and by confirming that the tier A-B evidence stratum replicates the pooled estimate (OR 0.452; Supplementary Table S3).

In the case studies, complement C3 and C5 anaphylatoxins—basic fragments of acidic precursors —represent the class of evidence invisible under precursor-centric reporting but emergent from a region-resolved reading. These fragments are established inflammatory biomarker species^14,20^. The detection of C3a (49 peptides, nine-study recurrence) despite the basic-class depletion is not a contradiction: an odds ratio of 0.45 describes a probabilistic reduction, not an absolute bar, and C3a circulates at concentrations high enough to overcome the analytical penalty. Even when the instrument measures a basic fragment, standard reporting files it under the acidic parent, hiding it from registry searches. Conversely, histone H4 tail-region and chemokine-region detection illustrate the forward direction: the detected material is basic but less so than the annotation implies. The AMBP null control (zero divergence) shows the contrast is a detection-coverage property, not a method artifact. Chemokines, a cytokine sub-family, are already included (CCL21, CXCL10; exploratory RQ3 chemokine-region OR 4.9, q = 6 × 10^−6^); interleukin- and TNF-type cytokines, many at very low abundance, would need enriched or targeted data.

Three practical implications follow. First, report observed regions: a gene-level label cannot say which species was measured (complement C3 evidence may derive predominantly from the basic C3a fragment), so residue range and region-level pI and molecular weight should accompany it. Second, digestion-aware design is a hypothesis: the signal concentrates in missed-cleavage peptides, so reduced-missed-cleavage protocols or complementary proteases are the natural test, but the DDA-specific deficit in synthetic peptides (OR 0.734; ETD 1.01) shows digestion is not the whole mechanism. Third, report the pK model and class thresholds: magnitude moved from 0.45 to 0.92–0.94 across calculators; direction did not.

This study demonstrates an analytical phenomenon in public proteomics datasets; it does not establish that current clinical diagnoses are incorrect or that a specific biomarker should be adopted or rejected. Changing clinical reporting practice would require targeted experimental validation.

Our approach differs from single-study peptide-centric reanalyses^5,6^ in quantifying physicochemical divergence across studies under a pre-specified data freeze, and from correlation-based proteoform-group inference^7^ in providing digestion-matched class-composition tests rather than co-variation patterns. The framework is a validated pipeline for divergence quantification, not a claim that every biomarker study misses basic proteoforms. Without orthogonal molecular-weight constraints or top-down validation^21,22^, region-supported calls define minimal sequence footprint regions, not intact proteoform boundaries—a distinction enforced by the five-level confidence taxonomy. That distinction is now measured against paired top-down proteoforms^21^ (Extended Data Fig. 5): detected regions recover top-down sequence space but do not delimit proteoform boundaries, so the 41 upgraded regions are partially validated. Where a canonical label conceals proteolytic fragments (here, C3a/C5a) or alternative splice products (ED Fig. 8 shows a splice-isoform case), reporting observed regions rather than the gene-level label is the direct remedy.

The observed bias is distinct from previously reported sample-preparation effects^10^: those act on protein abundance and extraction chemistry, whereas our digestion-matched design makes every protein its own control—abundance cancels, and the measured difference is in the class composition of what the assay detects from each protein’s theoretical tryptic space. The near-identical ORs across abundance strata (0.407 vs 0.405) confirm this separation.

All analyses were performed on a pre-specified, sha256-checksummed data freeze dated 2026-09-17, with hypotheses, thresholds, model specifications, and random seeds documented before any inferential test was run. A machine-readable extraction registry (28 entries) maps every figure to its source table. The complement-anaphylatoxin class did not survive multiple-testing correction (q = 0.112) and is reported as an exploratory observation pending larger cohorts.

The cohort spans seven plasma- or serum-only studies and three that include non-plasma material (FFPE intervertebral-disc tissue, FFPE colorectal-tumour tissue paired with plasma-derived extracellular vesicles, and cerebrospinal-fluid extracellular vesicles). This matrix mix is a generalizability feature rather than a confound: the digestion-matched background is constructed per study from that study’s own parent proteins, decoupling matrix composition from the class-comparison design, and every study individually shows an odds ratio below 1. The plasma- and serum-derived stratum (eight studies, incorporating the plasma-derived extracellular-vesicle arm of the paired colorectal cohort PXD069732) replicates the pooled estimate (OR 0.442), as do the DIA-only (OR 0.438, k = 4) and DDA-only (OR 0.479, k = 6) acquisition-mode strata (Supplementary Table S3). The between-study heterogeneity (I² = 97.6%) reflects variation in magnitude, not direction. Additional limitations include: the digestion-matched background assumes trypsin, and alternative proteases are untested; incomplete charge-state availability (three of ten studies export precursor charge); reliance on sequence-derived pI as a proxy for gas-phase behaviour; isoform ambiguity in protein-group assignments; and the absence of numeric molecular-weight constraint windows that would gate fragment-constrained calls.

## Methods

### Study selection and data acquisition

We screened 135 candidate datasets from the PRIDE Archive^26^ using 28 logged query families (disease-area, biofluid, DIA-acquisition, and tissue synonyms). Studies were scored on a written rubric (parser-family balance, file accessibility, sample-type fit, protocol-derived metadata quality, and a fractionation-evidence bonus), with protein-group-level-only exports excluded as evidence-ineligible. Ten studies were selected (PXD008583, PXD052666, PXD054594, PXD055218, PXD056620, PXD057799, PXD060933, PXD068982, PXD069732, PXD071549); four candidates were rejected with documented reasons. Every source file carries a SHA-256 checksum in the freeze manifest. The original five-study pilot’s rows were hash-verified as bit-identical after the five-study extension.

### Cohort size and power considerations

The pre-specified rule required a minimum of ten studies before inferential testing, motivated by the observation that DerSimonian-Laird between-study variance estimates are unstable below approximately ten clusters. The initial five-study pilot was locked to descriptive reporting only; unlocking the inferential phase required the cohort extension to k = 10 under the written selection rubric. Because the analysis plan, thresholds, and random seeds were fixed before the data freeze, the SHA-256 checksummed freeze serves as the pre-specification record: it demonstrates that the k ≥ 10 threshold, the primary odds-ratio model, and all sensitivity specifications were locked prior to any hypothesis test. At the observation level, the 70,020 evidence rows and 5,396 multi-peptide proteins yield overpowered primary tests (RQ1 p = 0.0005; RQ2 p ≈ 7 × 10^−30^); the constrained case is the RQ3 subclass analysis (complement-anaphylatoxin q = 0.112), reported as exploratory pending larger cohorts.

### Peptide evidence harmonization

Five parser families extract peptide-level evidence: MaxQuant peptides tables (*peptides.txt*, *peptides_Tissue.txt*), MaxQuant PSM tables (*msms.txt*), DIA-NN precursor reports (*DIA-NN_report.tsv*, *report.tsv*), Spectronaut facility peptide quantification (*FacilityPeptideQuant16.tsv*), Progenesis CSV (*peptides.csv*), and mzIdentML (via two-pass lxml reference resolution). Glycopeptide evidence from the per-run GlyPep-Quant outputs uses a naked-backbone policy: pI and molecular weight are computed on the unmodified peptide sequence, glycan composition is retained in ProForma-style notation, NeuAc/NeuGc-containing glycans are flagged as charge-altering, and glycopeptide rows are excluded from tryptic-background comparisons unless enzyme-matched. Every row carries a confidence tier (A: sequence + accession + q-value + intensity; B: sequence + accession + PEP/score; C: sequence + accession; D: no sequence, excluded from peptide-level analysis).

### UniProt mapping and join-quality gate

Protein-group strings are normalized (semicolon groups split, fasta-style prefixes stripped, isoform suffixes recorded) and resolved against UniProt^27^ via chunked REST stream queries (100 accessions per request, 0.6-s politeness interval, single retry) with a dated snapshot cache. Peptides are localized by exact substring matching; every row receives a five-tier mapping status. The join-quality gate operates as a four-stage programmatic firewall. First, raw protein-group strings are normalized: semicolon-delimited groups are split, FASTA-style prefixes (sp|/tr|) are stripped, REV_/CON_ decoy and contaminant tokens are flagged, and isoform suffixes (-2, -3) are logged before resolution. Second, accessions are resolved against UniProt via chunked REST stream queries (100 accessions per request, 0.6 s politeness interval, dated cache snapshot). Third, peptides are localized by exact substring matching against resolved canonical sequences; every row receives an explicit five-tier mapping status (exact-unique; exact multi-position; not found in group sequences; accession unmapped; no accession in source row) plus an evidence tier (A: sequence + accession + q-value + intensity; B: sequence + accession + PEP/score; C: sequence + accession; D: no sequence, excluded from peptide-level analysis). Fourth, the gate itself evaluates three pre-set thresholds: accession resolution at least 95% (observed 97.7%), exact-substring mapping at least 70% overall (observed 98.3% of 70,020 rows), and a per-study floor of at least 50% (observed minimum 90.8%, PXD056620, where 614 source-file rows lack protein assignments in the original msms.txt export). All three thresholds were cleared (gate: GO). Degenerate peptides and common contaminants (keratins, trypsins, serum albumin, immunoglobulin chains) are flagged per row rather than dropped.

### Physicochemical computation and validation

Theoretical pI and molecular weight are computed with Biopython (Bjellqvist pK set, primary). The engine was benchmarked against live ExPASy ProtParam on 16 sequences spanning full-length plasma proteins, UniProt-feature-anchored complement fragments, and synthetic acidic/basic controls (agreement within 0.005 pI units, one extreme-composition outlier documented). Molecular weight agrees with UniProt-declared values across all 2,602 proteins at a median deviation of −11.5 ppm (95th percentile of absolute deviation 32 ppm). Calculator sensitivity was quantified under three pK configurations (Biopython Bjellqvist, Rodwell/EMBOSS lineage, Lehninger); the 7.0–7.5-band class-flip rate is 74.5%, making calculator choice a named sensitivity analysis in all inferential models. The complete net-charge equations, residue-specific terminal pKa values for all three calculator models, molecular-weight conventions, digestion and background-construction parameters, region-stitching formalization, statistical model equations, and charge-state test statistics are provided in Supplementary Methods.

### Region and proteoform inference

Exactly mapped peptide spans are joined into detected regions when separated by ≤25 residues; region pI and molecular weight are computed from the actual precursor-slice sequence. Five confidence levels apply: *peptide-only* (single peptide, no region claim); *region-supported* (≥2 distinct peptides in one gap-bounded span); *fragment-constrained* (region-supported + numeric molecular-weight window consistency); *proteoform-group candidate* (co-varying peptide sets or ≥2 glycoforms on one backbone); *proteoform high-confidence* (orthogonal validation). Region claims define minimal sequence footprint regions, not intact proteoform boundaries. Molecular-weight constraint evidence was audited per study from protocol and file metadata; no pilot study exposes numeric windows, so fragment-constrained calls equal zero (recorded absence).

### Digestion-matched background construction

For each parent protein with at least one exactly mapped peptide, an *in silico* tryptic digest generates all theoretical peptides (cleavage after K or R unless followed by P; up to two missed cleavages; length 7–60 residues). Missed-cleavage counts are derived from the sequence itself (internal uncut K/R), not from file metadata, ensuring parity between detected and background classifications. The background shares the same parent proteins as the detected set, cancelling protein-level expression differences. The full background comprises 965,062 unique tryptic peptides.

### Statistical analysis

*RQ1 (divergence):* For each protein with ≥2 unique peptides, the median detected-peptide pI minus precursor pI was tested against a digestion-matched permutation null (2,000 resamples per protein at matched peptide counts; seed 42; two-sided empirical p). *RQ2 (class bias):* Per-study odds ratios for the basic class (detected vs background) with Woolf 95% CIs; inverse-variance fixed-effect summary; DerSimonian–Laird random-effects at k = 10 (pre-specified threshold); leave-one-study-out envelope; bootstrap 95% CIs (1,000 multinomial draws). *RQ3 (exploratory):* Fisher exact tests with Benjamini–Hochberg FDR (q < 0.05). *Covariates:* Stratified ORs by peptide-length quartile, missed-cleavage stratum (mc0/mc1/mc2+), and detected-intensity median split. *Calculator sensitivity:* All class comparisons repeated under three pK models. Software: Python 3.11, scipy 1.17, numpy, pandas. Random seed 42 throughout.

### Charge-state supplementary analysis

Precursor charge states are available for 33,821 of 70,020 rows (48.3%) from three studies whose result formats export charge annotations. Non-integer values (116 rows; 2.5, 3.5, 4.5 from median-of-PSM aggregation) are rounded to the nearest integer. Theoretical charge at assay pH ∼2.6 is estimated as count(K) + count(R) + count(H) + 1 (N-terminus). Three hypotheses were pre-stated: H-z1 (missed-cleavage → high charge; Fisher exact); H-z2 (basic-pI class → high charge; Fisher exact); H-z3 (detected mc2+ charge distribution shifted below background theoretical; permutation, B = 5,000). Results: H-z1 supported (mc1+ z ≥ 3: 80.0% vs mc0: 25.9%; OR 11.46); H-z2 not supported (basic 30.1% vs acidic 34.4%; opposite direction); H-z3 supported (detected 98.4% vs background 100.0%; p = 2 × 10^−4^).

### Orthogonal validation

Two validations were run against the unchanged pipeline under a metrics contract frozen on 2026-10-06, before any computation. (i) *Paired top-down ground truth.* The bottom-up arm of Sadeghi et al.^21^ (published source-data PSM table; 433,453 rows; the PXD077545 deposition contains the top-down arm only) was harmonized through the standard evidence schema with Probability standing in for the absent q-value column (validation-specific tier semantics), mapped by the exact-substring mapper (99.8% accession resolution; 99.4% exact mapping; 99.8% positional agreement with the search engine’s own coordinates), and subjected to region inference verbatim across the stitching-gap grid g ∈ {5, 10, 15, 20, 25, 30, 40}. Top-down proteoform spans were parsed from the deposited TDP tables (accession + first/last residue as coordinate authority; both proteoform-string dialects stripped; zero span-length mismatches; 3,671 deduplicated spans on 288 proteins). Concordance metrics, boundary-offset bootstrap, the span-permutation null, and the sweep-adequacy rule are specified in Supplementary Methods Module 7. (ii) *Digestion-free detectability.* Per-run MaxQuant peptide tables for ProteomeTools PXD010595^24^ (888 run files across four acquisition methods, 222 standard-DDA runs; 158 pools) yielded 214,336 pool-observed peptides as the denominator and 192,612 DDA-observed peptides as the numerator; class assignment used the canonical pI engine at the pre-specified thresholds, with the basic-vs-acidic odds ratio (Woolf 95% CI, unconditional 0.5 correction) and a one-sided Fisher test against the pre-specified directional prediction OR < 1. A dedicated machine-readable extraction registry covers all validation figures.

### Data freeze and reproducibility

All hypothesis-relevant inputs were frozen on 2026-09-17 under a sha256-checksummed manifest (13 files verified). Hypotheses, thresholds, model specifications, and random seeds were specified in the freeze document before any inferential test (a dated, SHA-256-manifested internal record; no external registry was used). A machine-readable extraction registry (28 entries; ED Fig. 8 registered separately) maps every figure to its source table. An exploratory probe (criteria set before the scan) classified peptides as canonical, shared or isoform-exclusive using UniProt isoforms. All notebooks execute end-to-end without manual intervention.

## Author contributions

S.S. conceived and designed the study, developed the reanalysis pipeline and software package, performed all analyses, and wrote the manuscript. V.A.-S. contributed key conceptual input to the study design and interpretation. M.R.B. and C.T. provided critical peer review that substantially strengthened the analysis and its orthogonal validation. G.M., S.N.B.N. and M.Y. reviewed and approved the final version. All authors read and approved the final manuscript.

## Competing interests

The authors declare no competing interests.

## Data availability

All source datasets are publicly available through the PRIDE Archive (accessions PXD008583, PXD052666, PXD054594, PXD055218, PXD056620, PXD057799, PXD060933, PXD068982, PXD069732, PXD071549). Processed peptide-evidence, mapping, feature, background, region, case-study, and charge-state tables, together with their SHA-256-checksummed freeze manifest, are provided as Supplementary Data. Analyses were performed on a versioned, checksummed data freeze dated 2026-09-17. The orthogonal-validation resources are also public: PXD010595 (ProteomeTools synthetic peptides; CC0) via PRIDE, and PXD077545 (top-down arm) via PRIDE, with the paired bottom-up PSM table published as the source data of Sadeghi et al.^21^ (SHA-256 recorded in the frozen validation contract).

## Code availability

The complete pipeline (harmonization, mapping, physicochemical computation, region inference, statistical modelling, case-study, charge-state, and figure-generation notebooks) is available at https://github.com/ShoabSaadat/proteoform-regions (public repository), installable directly from the Python Package Index (pypi.org/project/proteoform-regions; documentation at shoabsaadat.github.io/proteoform-regions). Each notebook executes end-to-end in the supplied Python environment. In addition, the method is distributed as the installable Python package **proteoform-regions** (command-line interface, documentation, worked walkthrough on an independent dataset, and a test suite whose freeze-parity checks reproduce the fixed-effect estimate reported here from the checksum-pinned inputs; source code archived on Zenodo, DOI 10.5281/zenodo.23214089, all-versions DOI 10.5281/zenodo.23214088), so the analysis can be re-run — or applied to new studies — without reconstructing the notebook environment.

## Extended Data

The following items are published in the online article PDF alongside the main figures. They do not count toward the six main display items.

**Extended Data Figure 1.**
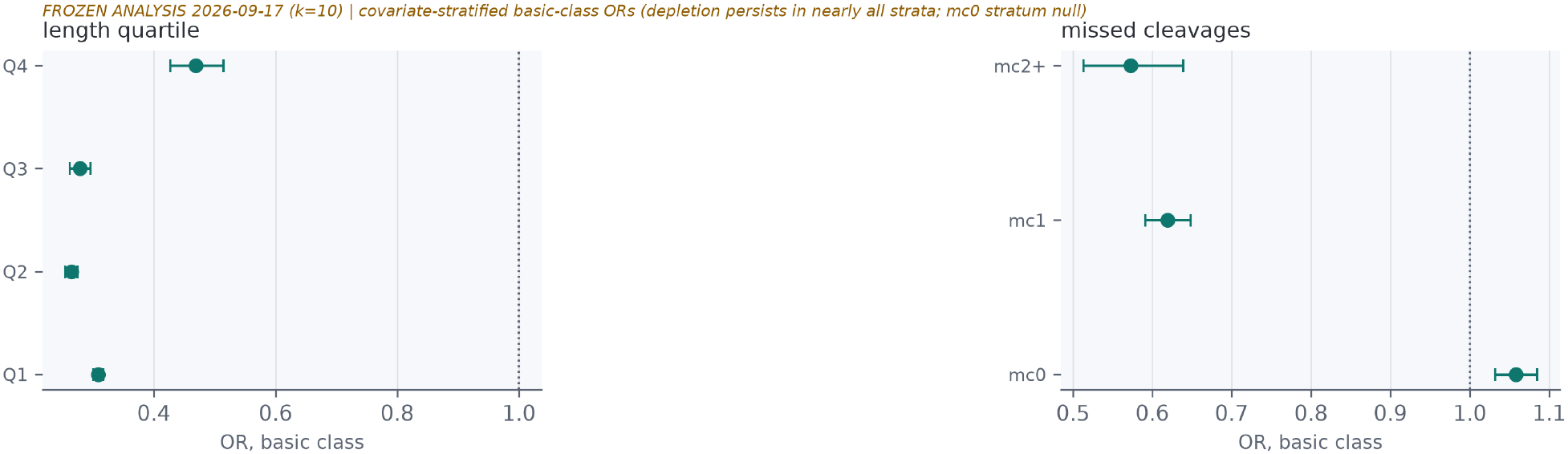
| Covariate-stratified basic-class odds ratios by missed-cleavage stratum and peptide-length quartile. The basic-class depletion persists across length quartiles (weakest in the longest, Q4). The fully-tryptic stratum (mc0) is null (OR 1.06), while missed-cleavage peptides deplete (mc1: 0.62; mc2+: 0.57), localizing the effect to digestion incompleteness.

**Extended Data Figure 2.**
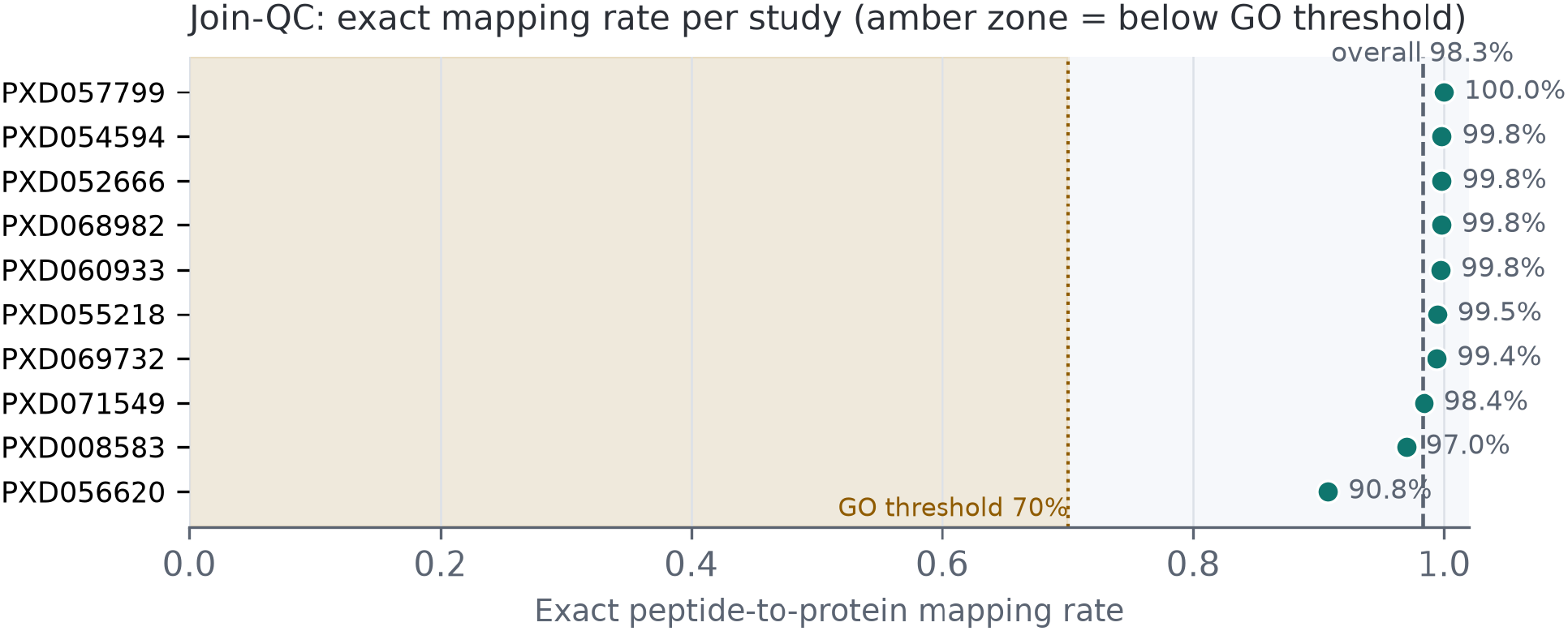
| Join-quality gate: per-study exact mapping rates. Teal points = per-study rates; dashed grey = overall reference; amber zone = below the 70% GO threshold. All studies pass.

**Extended Data Figure 3.**
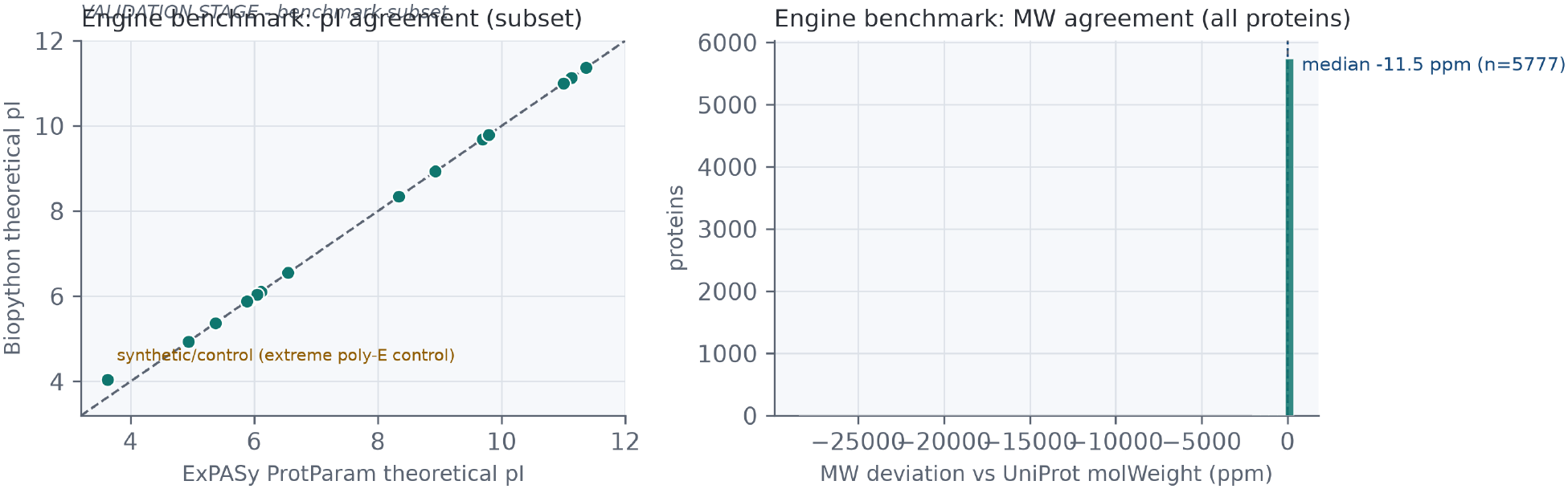
| Engine benchmark deviations. Left: pI agreement vs ExPASy ProtParam (identity line). Right: MW deviation vs UniProt molWeight (median −11.5 ppm).

**Extended Data Figure 4.**
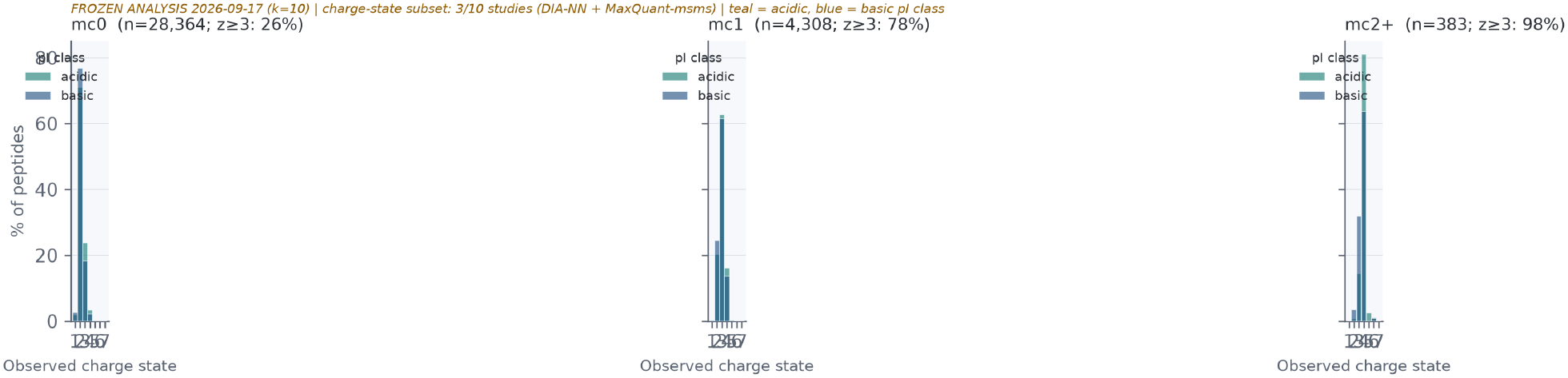
| Charge-state distributions by missed-cleavage stratum and pI class. Teal = acidic-class, blue = basic-class peptides. The mc1+/mc2+ strata show a pronounced shift toward z ≥ 3 (80.0%) vs mc0 (25.9%).

**Extended Data Figure 5.**
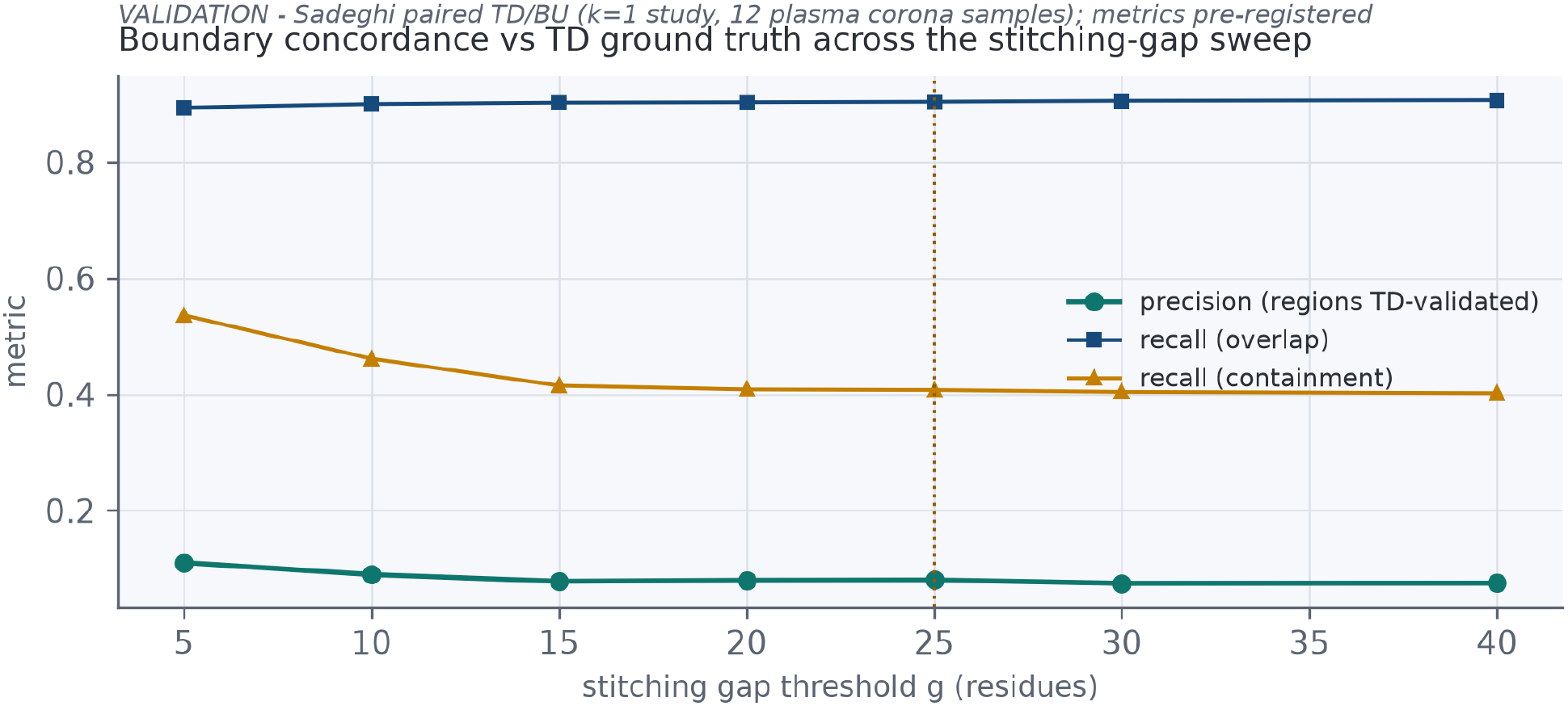
| Region-inference concordance vs paired top-down proteomics across the stitching-gap sweep. **Sadeghi paired TD/BU benchmark.** Precision (regions ≥80% contained in a top-down span, teal) and the recall family (blue/amber) across g = 5–40; dotted amber line marks the published g = 25. Precision stays at 7.5–11.0% while span-overlap recall stays ≥0.89: coverage is strong, boundary specificity is not. Metrics pre-specified; 237 evaluable proteins; 3,404 top-down spans.

**Extended Data Figure 6.**
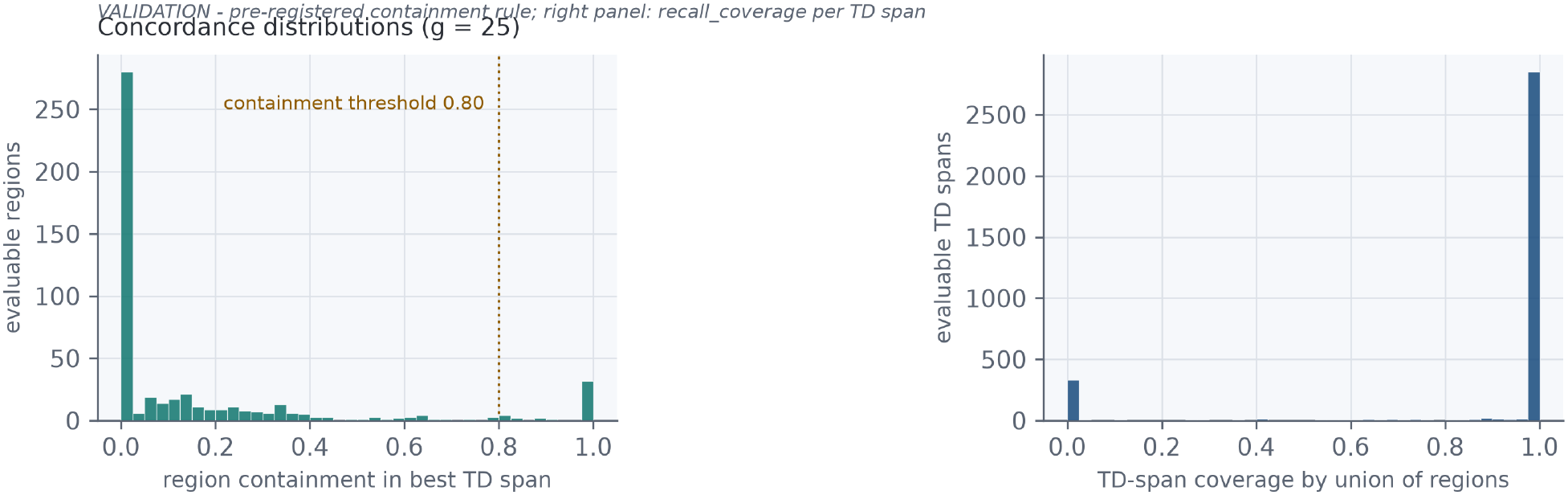
| Containment and coverage distributions at g = 25. Left: region containment in the best top-down span—most regions extend beyond any single proteoform (bulk near zero), with a secondary cluster at full containment (the 41 validated regions). Right: fraction of each top-down span covered by overlapping regions—median 1.00. Validated-region boundary offsets: start median 0 (CI 0–2), end median −2 (CI −5 to 0).

**Extended Data Figure 7.**
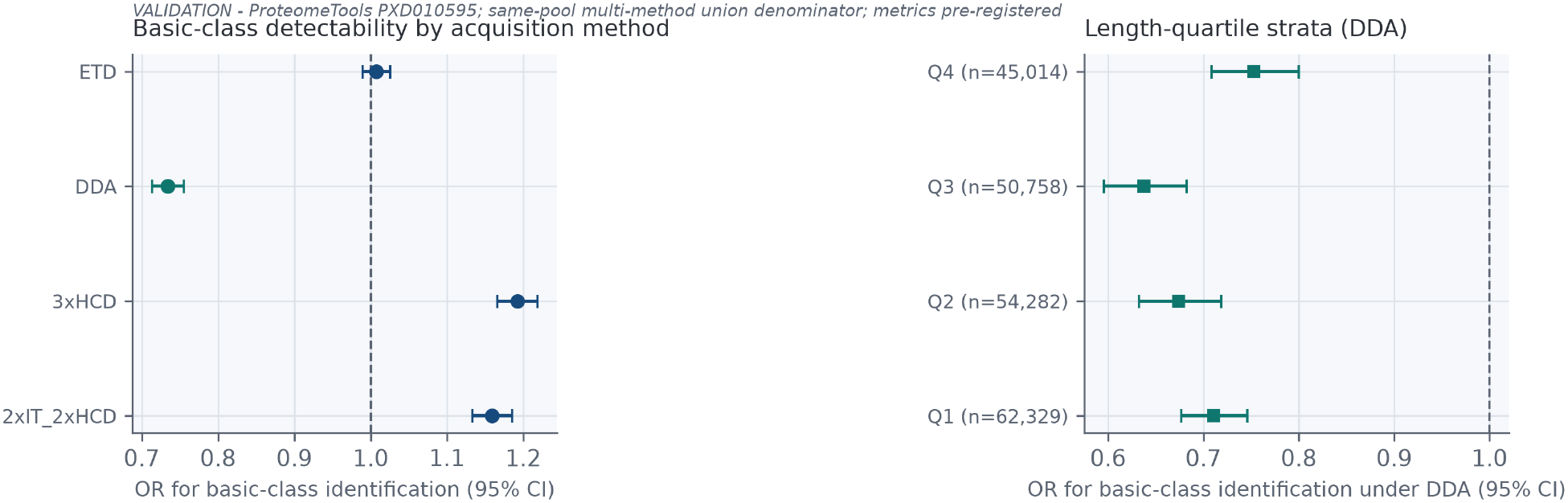
| Digestion-free basic-class detectability on synthetic peptides. **ProteomeTools PXD010595.** Left: odds ratios for basic-class identification per acquisition method against the same-pool multi-method union (DDA 0.734 highlighted; ETD 1.01; 2xIT-2xHCD 1.16; 3xHCD 1.19). Right: DDA odds ratios by peptide-length quartile (all <1). 214,336 peptides; metrics pre-specified.

**Extended Data Figure 8.**
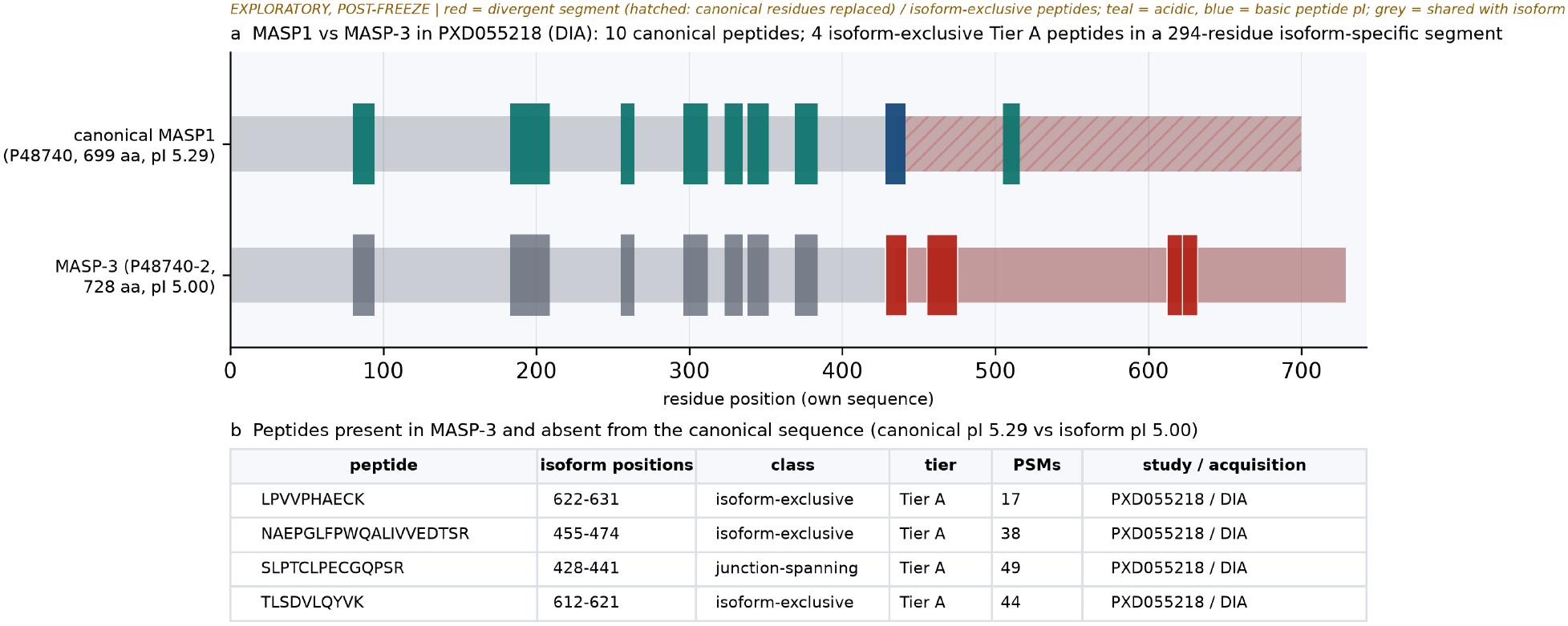
| Isoform evidence for MASP1 (exploratory, post-freeze). **MASP1 versus MASP-3 in PXD055218 (DIA).** a, Canonical MASP1 (P48740, 699 aa) and the annotated MASP-3 isoform (P48740-2, 728 aa). Ten canonical peptides are coloured by pI class (teal acidic, blue basic; grey where shared with the isoform); four Tier A isoform-exclusive peptides (red) fall in a 294-residue isoform-specific segment (hatched on the canonical sequence). b, Exclusive peptides with isoform positions and PSM counts; canonical pI 5.29, isoform pI 5.00. Exploratory, post-freeze; one study, no replication; no statistic is claimed.

## Supplementary Methods: Mathematical and Algorithmic Specifications

### Module 1: Physicochemical engine

#### Isoelectric point (pI)

The primary pI calculator is Biopython’s `IsoelectricPoint` module (version 1.88), which computes the theoretical pI by finding the pH at which the net charge Q(pH) = 0. The net charge is the sum of positive contributions from the N-terminus and the side chains of Lys, Arg, and His, minus negative contributions from the C-terminus and the side chains of Asp, Glu, Cys, and Tyr:

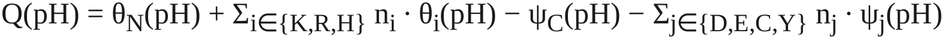

where the partial charge of a basic group with pK_b_ is θ(pH) = 1/(1 + 10^pH−pKb^), and the partial charge of an acidic group with pK_a_ is ψ(pH) = 1/(1 + 10^pKa−pH^). A key feature of the Biopython implementation is that the N-terminal and C-terminal pK values are *residue-specific*: the N-terminal pK depends on the identity of the first residue (A = 7.59, M = 7.0, S = 6.93, P = 8.36, T = 6.82, V = 7.44, E = 7.7; default 7.5), and the C-terminal pK depends on the last residue (D = 4.55, E = 4.75; default 3.55).

#### Sensitivity calculators

Two alternative pI models were implemented as custom Henderson– Hasselbalch charge functions using pK tables from the pyteomics.electrochem module, solved by 60-iteration bisection between pH 0 and 14 (convergence is implicit; precision ≈ 14/2^60^).

**pK lookup table (side chains and termini):**

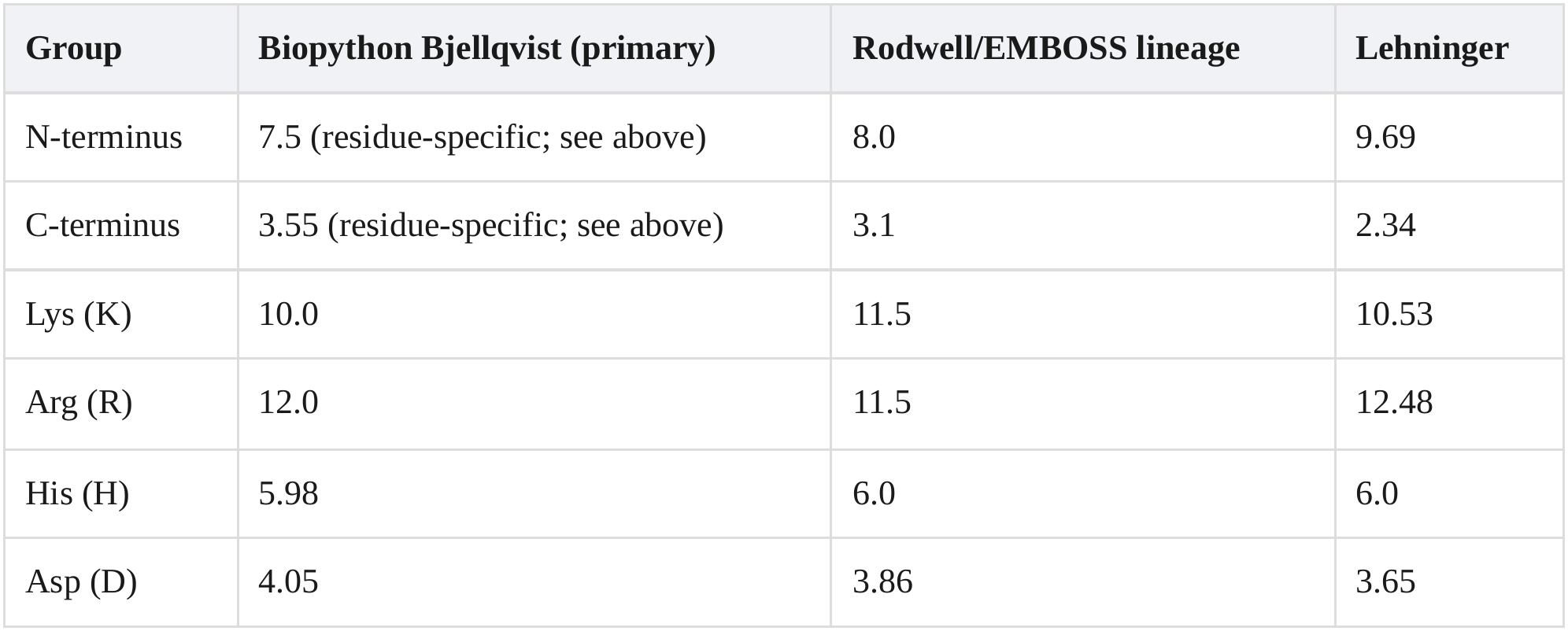
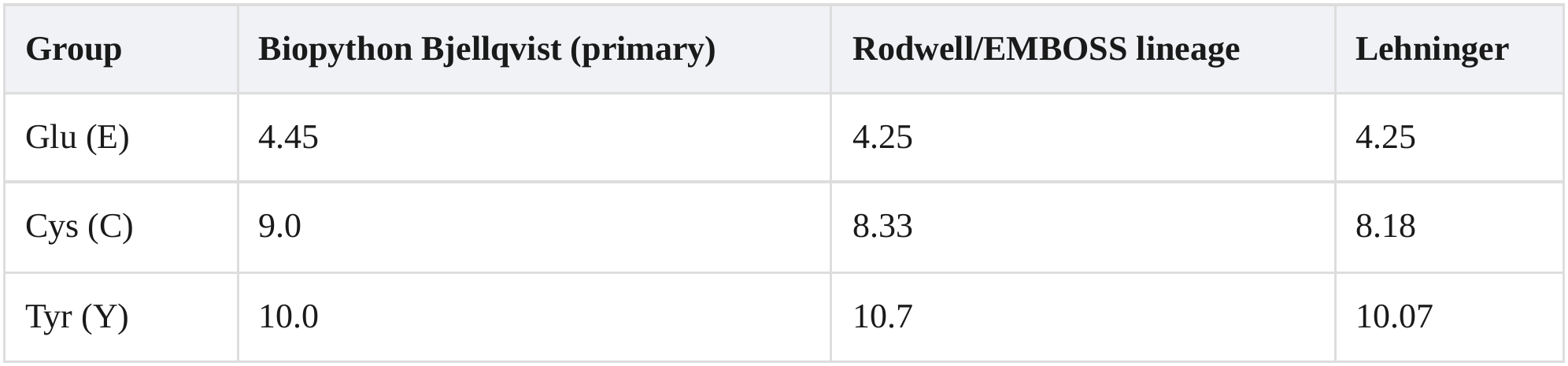

#### Molecular weight

MW is computed via Biopython’s `ProteinAnalysis.molecular_weight()`, which uses the *average isotopic mass* of each amino acid residue plus terminal water (H_2_O = 18.015 Da). Non-canonical residue X is replaced by alanine before all physicochemical calculations. No chemical modifications (carbamidomethylation, oxidation) are applied to the theoretical backbone, matching the unmodified nature of the in silico background.

#### Benchmark methodology

The primary engine was validated against (i) live ExPASy ProtParam on 16 curated sequences (7 full-length plasma proteins, 4 UniProt-feature-anchored complement fragments, 5 synthetic acidic/basic controls), yielding agreement within 0.005 pI units except one extreme-composition poly-E control; and (ii) UniProt-declared molWeight across all resolved proteins (median deviation −11.5 ppm; 95th percentile of absolute deviation 32 ppm).

### Module 2: In silico digestion and digestion-matched background construction

#### Cleavage rule

Theoretical tryptic peptides are generated per parent protein by applying the Keil rule: cleave after lysine (K) or arginine (R) unless the subsequent residue is proline (P). Missed cleavages are enumerated as 0, 1, or 2 internal uncut K|R sites (counted from the sequence itself, not from file metadata). The peptide length filter is 7 ≤ L ≤ 60 residues.

#### Deduplication scope

Background peptides are deduplicated per (accession, peptide-sequence) pair — that is, within each parent protein. The same peptide sequence arising from two different parent proteins is retained as two separate background entries, preserving the per-protein matching that underlies the digestion-matched design.

#### Initiator methionine

No special handling: digestion starts from position 1 of the canonical UniProt sequence. Initiator-Met excision is not simulated.

#### Non-canonical residues

Residue X is replaced by alanine (A) before pI and MW calculations. Selenocysteine (U) and pyrrolysine (O) are not present in the charged-residue pK tables and are therefore effectively ignored by the charge calculation (they contribute 0 to Q(pH)).

#### Background scope

For each study, the background comprises the theoretical tryptic peptides of all parent proteins that have at least one exactly mapped detected peptide in that study. This per-study construction ensures the background and the detected set share the same precursors, cancelling protein-level expression differences. The full background across the 10-study cohort comprises 965,062 unique (accession, peptide) pairs.

### Module 3: Region-stitching algorithm

#### Single-linkage clustering

For each (study, protein) pair, exactly mapped peptide spans [s_i_, e_i_] (1-based, inclusive) are sorted by start position. A greedy single-linkage merge proceeds left to right: consecutive spans are joined into the same cluster when the next span’s start position satisfies s_j_ − e_i_ ≤ 25 (where e_i_ is the end of the previous span in the current cluster). This means the maximum start-to-end distance between adjacent mapped spans is 25 residues (equivalently, at most 24 unobserved intervening residues).

#### Sequence slice

For each cluster, the detected region R_k_ is defined as the continuous sequence slice P[s : e] from the canonical precursor, where s = min(s_i_) and e = max(e_j_) across all spans in the cluster. This slice *includes* the unobserved intermediate residues between mapped peptides. Region pI and MW are computed on this actual precursor-slice sequence (not as an average of member-peptide values).

#### Minimum evidence

A region is classified as *region-supported* if it contains ≥ 2 distinct peptide sequences. Single-peptide clusters remain *peptide-only*.

#### Terminal-inclusive flag

A region is flagged N-terminal-inclusive if s ≤ 0.10 × L (precursor length), and C-terminal-inclusive if e ≥ 0.90 × L.

### Module 4: RQ1 permutation null model

#### Per-protein null

For protein *i* with *n_i_* observed unique peptides, let T_i_ be the set of theoretical tryptic peptide pI values from the digestion-matched background of that protein. In each of B = 2,000 permutation iterations (seed = 42), a random sample of size *n_i_* is drawn *with replacement* from T_i_ (i.e., a multinomial bootstrap), and the median of the sampled pI values minus the precursor pI is recorded as ΔpI_null,i,b_.

#### Edge case

Proteins whose background contains fewer distinct peptides than the detected count (len(T_i_) < *n_i_*) are skipped (n = 5,388 of 5,396 pass; 8 skipped).

#### Per-protein empirical p

p_i_ = (1 + Σ_b_ 1[|ΔpI_null,i,b_| ≥ |ΔpI_obs,i_|]) / (B + 1). The minimum achievable p is 1/2001 ≈ 0.0005.

#### Global test

The global statistic T_obs_ is the median of the observed per-protein divergences. The global null distribution is constructed by resampling (with replacement) the set of per-protein null-divergence medians B_boot_ = 2,000 times and taking the median of each resample. The global empirical p-value is: p = (1 + Σ_b_ 1[|median(null_b_) − median(null)| ≥ |T_obs_ − median(null)|]) / (B_boot_ + 1).

### Module 5: Meta-analytic models (RQ2)

#### 2 × 2 contingency table

For each study, unique detected peptides and unique background peptides are classified as basic (pI ≥ 7.5) or non-basic (pI < 7.5; includes both acidic and near-neutral):

|  | **Basic** | **Non-basic** |
|---|---|---|
| **Detected** | a | b |
| **Background** | c | d |

#### Odds ratio and Woolf SE

OR = (a·d)/(b·c); SE(ln OR) = √(1/a + 1/b + 1/c + 1/d). A continuity correction of max(cell, 0.5) is applied *unconditionally* to all four cells (i.e., a Haldane–Anscombe correction applied always, not only when a cell is zero).

#### Fixed-effect summary (inverse variance)

w_i_ = 1/SE_i_²; ln OR_FE_ = Σw_i_·ln OR_i_ / Σw_i_; SE_FE_ = √(1/ Σw_i_); 95% CI = ln OR_FE_ ± 1.96·SE_FE_.

#### DerSimonian–Laird random-effects

Q = Σw_i_(ln OR_i_ − ln OR_FE_)²; τ² = max(0, (Q − (k−1)) / (Σw_i_ − Σw_i_²/Σw_i_)); w_i_* = 1/(SE_i_² + τ²); ln OR_RE_ = Σw_i_*·ln OR_i_ / Σw_i_*. I² = max(0, (Q − (k−1))/Q) × 100%.

#### Leave-one-study-out envelope

The fixed-effect summary is recomputed excluding each study in turn; the LOO envelope is the range of the 10 resulting summary ORs.

#### Bootstrap CIs (descriptive layer)

For the descriptive enrichment ratios (detected share / background share per class), 95% percentile CIs are computed via 1,000 multinomial draws from the observed class-count vector independently for detected and background sets (seed = 42). The inferential ORs in S8 use the Woolf normal approximation rather than the bootstrap.

### Module 6: RQ3 and charge-state statistics

#### RQ3 Fisher exact tests

For each disease-relevant region class (complement_anaphylatoxin, histone_gene, chemokine_gene, histone_tail_candidate), a 2 × 2 contingency table is constructed over all detected regions:

|  | **Basic (pI ≥ 7.5)** | **Non-basic** |
|---|---|---|
| **Flagged class** | n<sub>flag,basic</sub> | n<sub>flag,non-basic</sub> |
| **All other regions** | n<sub>unflag,basic</sub> | n<sub>unflag,non-basic</sub> |

Fisher’s exact test (two-sided) is computed; the Benjamini–Hochberg correction is applied across the m = 4 hypotheses in this family. The BH adjusted q-value for the i-th ranked p-value is q_(i)_ = min_k≥i_ {m·p_(k)_/k}, capped at 1.

#### Charge-state H-z1 and H-z2 (Fisher exact)

The same 2 × 2 table structure is used with rows = stratum (mc0 vs mc1+ for H-z1; basic-pI vs acidic-pI for H-z2) and columns = charge state (z ≥ 3 vs z < 3).

#### Charge-state H-z3 (one-sided permutation)

The test statistic is the proportion of detected mc2+ peptides with observed charge z ≥ 3. Under the null, the background mc2+ theoretical-charge distribution is resampled with replacement at the detected-set size B = 5,000 times (seed = 42), and the proportion with z ≥ 3 is computed for each resample. The one-sided p-value is: p = (1 + Σ_b_ 1[null share_b_ ≤ observed share]) / (B + 1).

### Module 7: Orthogonal-validation metrics (Barnes round)

#### Frozen contract

All metrics below were pre-specified and frozen (2026-10-06, sha256-verified inputs) before any computation; two recorded errata (digest transcription; sanity-direction wording) affected no metric definition.

#### Evaluable frame (paired TD/BU)

P_TD_ = canonical accessions with ≥1 top-down span in the primary union (TDP_CZE_ ∪ TDP_LC_, deduplicated by accession + first + last residue + mass at 2 dp); P_BU_ = accessions with ≥1 exactly-mapped bottom-up peptide; the evaluable set is P_TD_ ∩ P_BU_. For region R = [r_s_, r_e_] and top-down span T = [t_s_, t_e_] on the same protein: overlap(R,T) = max(0, min(r_e_, t_e_) − max(r_s_, t_s_) + 1), containment_R_ = overlap/|R|, Jaccard(R,T) = overlap/(|R| + |T| − overlap)

The best pair for a region maximizes Jaccard (ties broken by larger overlap).

#### Endpoints per stitching gap g ∈ {5, 10, 15, 20, 25, 30, 40}

A region is TD-validated iff max_T_ containment_R_ ≥ 0.80. Precision = validated evaluable regions / evaluable `region_supported` regions; recall_overlap_ = evaluable spans with overlap ≥ 1 residue / evaluable spans; recall_coverage_ = median over evaluable spans of the union-covered fraction of the span; recall_containment_ = fraction of spans with a region where containment_R_ ≥ 0.80 and overlap/|T| ≥ 0.20. Boundary offsets on best pairs of validated regions: Δ_s_ = r_s_ − t_s_, Δ_e_ = r_e_ − t_e_; 95% CIs by protein-level bootstrap (B = 2,000, percentile, seed 42). Permutation null: TD span lists shuffled across proteins within ±20% precursor-length bins (B = 5,000, seed 42); statistic median|Δ_s_| + median|Δ_e_|; p = P(null ≥ observed) as frozen.

#### Adequacy rule

g = 25 is retained iff precision(25) ≥ 0.70 and precision(25) ≥ max_g_ precision(g) − 0.05; otherwise the precision-maximizing g is reported and the 25-residue choice is flagged for re-examination.

#### Digestion-free detectability (PXD010595)

Denominator = union of peptides observed by any of the four acquisition methods within DDA-covered pools; numerator = peptides observed in ≥1 of the 222 standard-DDA runs. Class assignment by the canonical pI engine (acidic < 7.0 ≤ near-neutral < 7.5 ≤ basic; near-neutral excluded from the 2 × 2). OR = (a·d)/(b·c), with a, b, c, d Woolf-corrected (unconditional 0.5); SE(ln OR) = √(1/a + 1/b + 1/c + 1/d), with a one-sided Fisher exact test against the pre-specified directional prediction OR < 1; sensitivity by acquisition method, peptide-length quartile, and the three pK models.

## Supplementary Note 1: Provenance and stage-wise validation

### Stage 1: Discovery and harmonization

The 135-study screen produced a scored shortlist; the initial five-study pilot tranche was extracted through the five parser families (23,957 rows), with a preview-guard system ensuring that schema-probe rows never entered statistical computation. The extension to ten studies used a written selection rubric with hash-verified pilot-row immutability (Supplementary Fig. S1).

### Stage 2: UniProt mapping

Accession normalization and exact substring mapping achieved 97.7% resolution and 98.3% exact mapping at the extended k=10 cohort. A pre-thresholded join-quality gate (≥95% resolution, ≥70% mapping, ≥50% per study) passed (Extended Data Fig. 2).

### Stage 3: Physicochemical engine validation

The Biopython Bjellqvist engine was benchmarked against live ExPASy ProtParam on curated standards and UniProt-declared molecular weights (Extended Data Fig. 3). A three-calculator class-flip matrix quantified the 7.0–7.5-band sensitivity that motivates the named sensitivity analysis (Supplementary Table S3).

### Stage 4: Region inference

Detected regions were constructed by gap-bounded stitching (≤25 residues) of exactly mapped peptide spans, yielding 20,515 regions with five-level confidence labels (Supplementary Fig. S2). No pilot study provided numeric molecular-weight constraint windows; this absence was documented rather than bypassed (Supplementary Fig. S3).

### Stage 5: Cohort extension

Five additional studies were selected by rubric, downloaded with SHA-256 manifests, and extracted through the existing parser architecture. The evidence table grew from 23,957 to 70,020 rows; the original pilot rows were verified bit-identical after append.

### Stage 6: Data freeze

All hypothesis-relevant inputs were checksummed and frozen on 2026-09-17 (13 files, 13/13 sha256 verified). Hypotheses, thresholds, model specifications, and seeds were specified in the freeze document before any inferential test.

### Stage 7: Inference

Permutation tests, odds ratios, and meta-analytic summaries were computed on the frozen tables with pre-stated rubrics (gate G1: answer (a), proceed). Covariate stratification, calculator sensitivity, and leave-one-study-out envelopes were executed as pre-specified (Extended Data Fig. 1).

### Stage 8: Case studies

Six cases were selected by divergence magnitude, disease-class annotation, peptide count, and cross-study recurrence, anchored by a methodological null control (Fig. 4).

## Supplementary Information index

**Supplementary Note 1.** Provenance and stage-wise validation (eight stages from discovery through case studies; see above).

**Supplementary Figure S1.** Preview-vs-full physicochemical comparison across parser families.

**Supplementary Figure S2.** Detected-region span and coverage distributions by study.

**Supplementary Figure S3.** MW-constraint evidence availability per study.

**Supplementary Figure S4.** Disease-relevant class-flag counts on detected regions.

**Supplementary Figure S5.** Abundance vs pI hexbin density (n = 52,848 rows with intensity values).

**Supplementary Table S1.** Per-study extraction summary.

**Supplementary Table S2.** Tier-confidence crosswalk.

**Supplementary Table S3.** Full calculator-sensitivity results.

**Supplementary Table S4.** Case-study detail.

**Supplementary Table S5.** Charge-state analysis results.

**Supplementary Methods.** Modules 1–6 (engine, digestion/background, region stitching, RQ1 null, meta-analysis, RQ3/charge-state) and Module 7 (orthogonal-validation metrics).

**Source Data.** One file per main and Extended Data figure.

## Notes

### Competing Interest Statement

The authors have declared no competing interest.

